# Acute Neuropixels recordings in the marmoset monkey

**DOI:** 10.1101/2023.12.14.571771

**Authors:** Nicholas M. Dotson, Zachary W. Davis, Patrick Jendritza, John H. Reynolds

## Abstract

High-density linear probes, like Neuropixels, provide an unprecedented opportunity to understand how neural populations within specific laminar compartments contribute to behavior. Marmoset monkeys, unlike macaque monkeys, have a lissencephalic (smooth) cortex that enables recording perpendicular to the cortical surface, thus making them an ideal animal model for studying laminar computations. Here we present a method for acute Neuropixels recordings in the common marmoset (*Callithrix jacchus*). The approach replaces the native dura with an artificial silicon-based dura that grants visual access to the cortical surface, which is helpful in avoiding blood vessels, ensures perpendicular penetrations, and could be used in conjunction with optical imaging or optogenetic techniques. The chamber housing the artificial dura is simple to maintain with minimal risk of infection and could be combined with semi-chronic microdrives and wireless recording hardware. This technique enables repeated acute penetrations over a period of several months. With occasional removal of tissue growth on the pial surface, recordings can be performed for a year or more. The approach is fully compatible with Neuropixels probes, enabling the recording of hundreds of single neurons distributed throughout the cortical column.

## INTRODUCTION

Recent advancements in electrophysiological recording tools have made it possible to study cortical function at unprecedented scales, including simultaneous recordings of hundreds to thousands of neurons across the depth of cortical layers using Neuropixels probes (Steinmetz et. al., 2021; Siegle et. al., 2021). In the macaque monkey, many of these tools have limited utility for studying laminar processing because of the extensive gyrification of the neocortex, making it challenging in many brain areas to reliably obtain perpendicular penetrations of the cortex with linear electrode arrays (Figure 1). The marmoset has been gaining interest as a model organism for studying cortical function due to its lissencephalic (smooth) cortex and its ability to perform complex visual tasks (Mitchell et. al., 2014; Solomon & Rosa, 2014; Selvanayagam et. al., 2019; Davis et. al., 2020; Cloherty et. al., 2020; Feizpour et. al., 2021; Jendritza et. al., 2021; Kell et. al., 2023; Jendritza et. al., 2023; Piza et. al., 2023; Davis et. al., 2023; Dotson et. al., 2023). However, there are still numerous technical considerations that can pose challenges to maximizing the utility of the marmoset cortex for neuroscience research. The marmoset skull, compared to that of a macaque, is small and thin (∼1 mm thick) and therefore requires the use of lightweight materials and small parts. Similar to the macaque monkey, the marmoset dura is opaque and difficult to penetrate with laminar probes, often requiring its removal. However, removing the dura in order to access the brain introduces risks of infection or damage and results in the formation of coarse granulation tissue that must be periodically removed (Sadakane et al. 2015).

**Figure 1:**
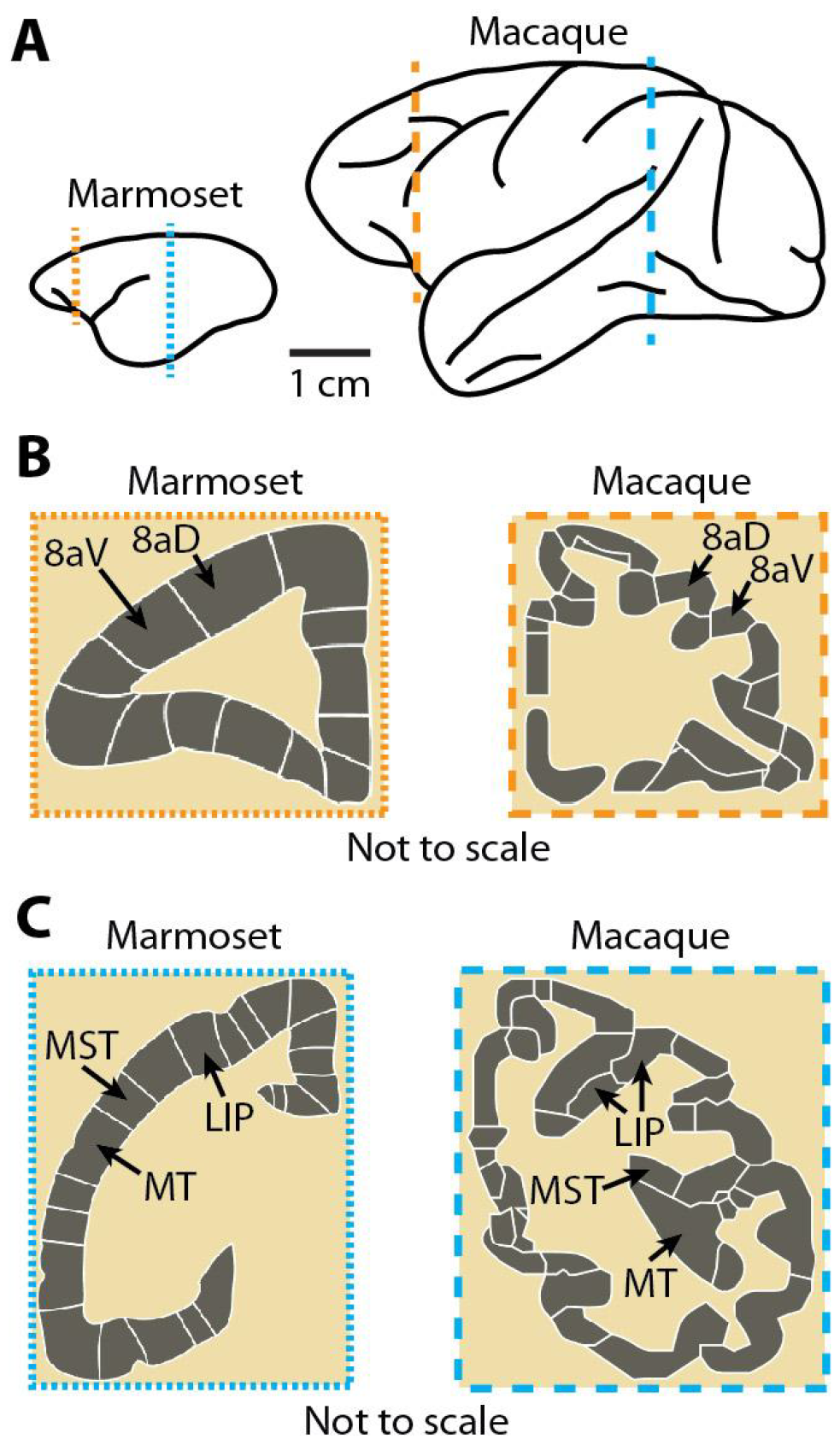
The near absence of sulci in the marmoset makes the cortex highly suitable for laminar recordings. A) Trace drawings of the marmoset and macaque monkey brains illustrating their relative sizes and the differences in the prevalence of sulci. The marmoset cortex is mostly smooth, while the macaque cortex is highly convoluted making it difficult to record perpendicular to the surface in many locations of the cortex. B-C) Digital reconstructions (not to scale relative to each other) of coronal slices further illustrate the relative ease of making laminar recordings in the marmoset compared to the difficulties with performing laminar recordings in the macaque monkey. Cortical areas 8aV, 8aD, MT, MST, and LIP are highlighted to show the differences in position created by sulci. (Images in B and C are from the scalablebrainatlas.incf.org; Paxinos et. al., 2000; Van Essen, 2002; Tokuno et. al., 2009; Paxinos et. al., 2012; Bakker et. al., 2015)

Here we describe a technique for repeated acute recordings of laminar cortical activity using one or two Neuropixels probes (short, linear and sharpened NHP version) in the marmoset neocortex. Our method utilizes an artificial dura system similar to ones previously used in macaque monkeys (Shtoyerman et. al., 2000; Arieli et. al., 2002; Chen et. al., 2002; Gray et. al., 2007; Ruiz et. al., 2013; Nassi et. al., 2015; Nandy et. al., 2017; Franken & Reynolds, 2021), but modified to suit the specific challenges that arise due to the smaller size of the marmoset. The recording chamber provides mounts for attaching a microdrive and houses a removable insert that is fitted with an artificial dura (AD). The AD is made from transparent silicone elastomer, providing optical clarity of the cortical surface, thus permitting the insertion of Neuropixels and other high-density linear electrode arrays perpendicular to the cortical surface while avoiding blood vessels. The chamber is sealed using a silicone gasket which greatly reduces the risk of infection and can be flexibly designed to accommodate a variety of experimental aims, including other recording and stimulation techniques, such as optical imaging, optogenetics, and semi-chronic microdrives.

## RESULTS

### Artificial dura (AD) recording system for the marmoset

The artificial dura (AD) system consists of a chamber, an insert with the artificial dura adhered to the bottom, and a cap with a silicone gasket (Figure 2). The chambers are made of titanium, and multiple designs have been implemented to record either in area MT (Figure 2A) or in the prefrontal cortex (PFC) (Figure 2B). The maximum inner dimension of the chamber, not including the insert, ranges from 6 to 9.5 mm depending on the chamber design, providing access to multiple cortical areas. The chamber height is (3.1 mm) and the housing cap (1.6 mm) is secured with 4 screws placed strategically around the chamber perimeter. The shape of the chamber is specifically customized to fit the constraints of the implantation site. For example, the MT chamber (Figure 2A) has the screw holes placed more dorsally to reduce the impact on the ventral aspect of the skull near the marmoset ears. The PFC chamber (Figure 2B) was shaped to provide access to the more anterior cortical regions while reducing the implant’s footprint near the orbitofrontal bone and eyes. A thin skirt that extends beyond the bottom of the chamber passes into the craniotomy ensuring a tight fit.

**Figure 2:**
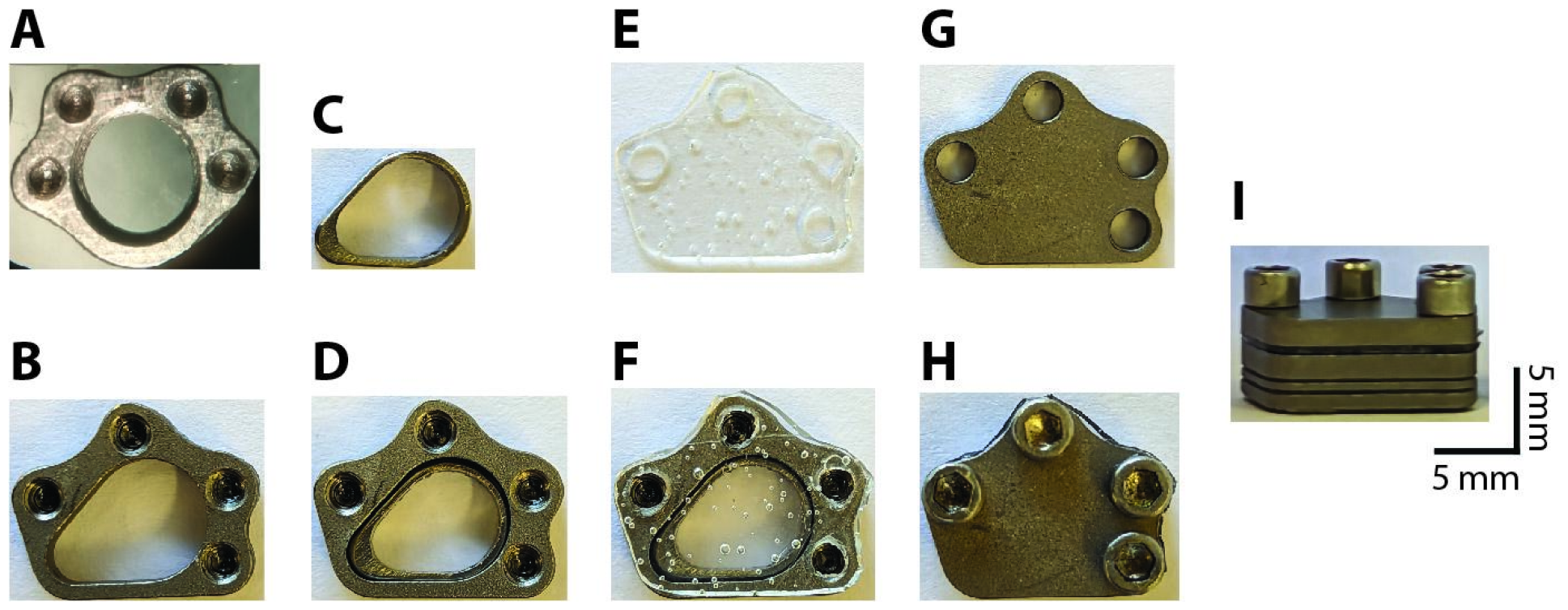
Recording chamber components. A) Image of circular recording chamber used for MT recordings (top view). B) Image of egg-shaped recording chamber used for PFC recordings (top view). C) Insert for PFC chamber (AD not attached). D) Chamber with the insert inside. E) Image of the silicone gasket used to seal the chamber. F) Image of the gasket on top of the chamber. G) Image of the cap for the PFC chamber. H) Image of the chamber with gasket, cap, and four screws fixing the cap in place (top view). I) Side view of the PFC chamber with gasket and cap. The scale bars on part (I) apply to all images.

The insert with the AD attached to the bottom fits tightly into the chamber and can be easily replaced, e.g. in the event that the AD tears or breaks (Figure 2C, D). The walls of the insert are 0.5-1 mm thick and can be made in various heights (e.g. 2.5 to 5 mm). This permits swapping out inserts to accommodate different distances to the cortical surface that may arise after surgical implantation of the chamber. The chamber and insert can be designed with a lip to restrict the maximum depth of the insert with respect to the bottom surface of the chamber (Figure 2A and Figure 3H).

**Figure 3:**
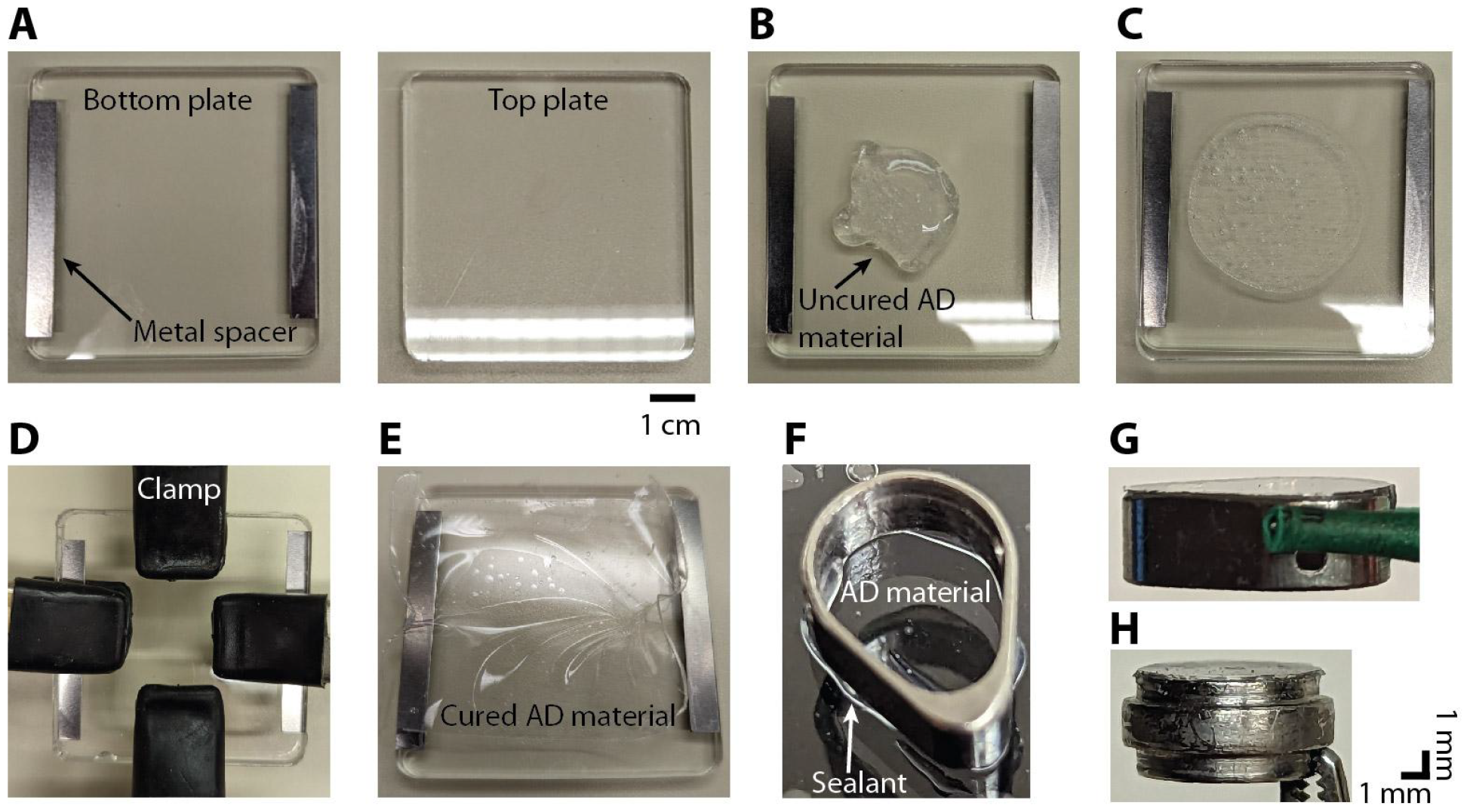
Procedure for making the AD material and adhering it to the insert. A) Two acrylic plates are used to cure the AD material into a thin sheet. Metal spacers are attached to the bottom plate (left) in order to create a gap of 400 μm between the two plates. B) Several milliliters of the uncured AD material (Shin-Etsu, KE-1300T) is applied to the bottom plate. C) The top plate is then pressed down on top of the bottom plate. D) Four hand clamps are used to secure the top plate to the bottom plate. To achieve a thinner AD thickness the clamps are placed towards the middle. This creates a thickness of 100 to 200 μm in the center. E) After 24 hours the cured AD material is in a thin sheet of the desired thickness. F) A thin bead of silicone sealant (DOWSIL, 734 flowable sealant) is applied to the bottom surface of the insert before gently pressing it down onto the AD material. G) Insert with AD attached. H) Image of the circular insert with AD attached. The bulging section in the middle of the round insert rests on a lip inside the round chamber. In G and H, the insert is oriented upside down with the AD material at the top.

The chamber is sealed between recordings by a 0.5 mm thick silicone gasket which is placed on top of the chamber followed by a titanium cap that is screwed down to provide an airtight seal (Figure 2E-I). This gasket is removed every time the chamber is opened for cleaning or recording and replaced with another sterile gasket and cap. The chamber is cleaned every 5 days or less by rinsing with sterile saline. The chamber is also rinsed with sterile saline before and after recording. Using this approach, we have not observed any evidence of infection in the four marmosets that have been implanted with this system.

The AD recording system is based on a previous design used in macaque recordings (Ruiz et. al., 2013; Nassi et. al., 2015; Nandy et. al., 2017; Franken & Reynolds, 2021). A 10:1 mixture of silicone to activator (Shin-Etsu, KE-1300T) is placed in a mold, fixed with clamps, and left to cure for 24 hours (Figure 3A-E). The bottom surface of the insert (Figure 2C) is slightly roughened using a file and then soaked in acetone to remove any debris or oils prior to attaching the AD material. A thin bead of silicone sealant (DOWSIL, 734 flowable sealant) is applied to the bottom surface of the insert. The insert is then pressed gently onto the AD which has been slightly stretched and placed on a flat surface (Figure 3F). After 24 hours a scalpel is used to cut around the perimeter of the insert. Any excess AD that protrudes over the edge of the insert is trimmed (Figure 3G, H). After autoclaving the insert is ready for use.

Recording chambers were implanted over area MT in two marmosets and over PFC in another two marmosets. Each animal was implanted with a headpost to stabilize the head for neurophysiological recordings and eye tracking. All surgical procedures were performed with the animal under general anesthesia in an aseptic environment in accordance with the recommendations in the Guide for the Care and Use of Laboratory Animals of the National Institutes of Health. All experimental methods were approved by the Institutional Animal Care and Use Committee (IACUC) of the Salk Institute for Biological Studies and conformed with NIH guidelines. Laminar probe recordings were performed in all animals, but Neuropixels recordings were performed only in the PFC marmosets.

### Single and multi-Neuropixels acute recording methods

Prior to recording, a custom 3D-printed holder is glued (Gorilla glue, 2 part epoxy) to the base plate of the Neuropixels probe (Figure 4A). A rod (1.13 mm) is inserted into the holder and glued in place (Gorilla glue, 2 part epoxy). Using standard parts, the probe can then be mounted to a microdrive (Narishige, MO-97A oil hydraulic micromanipulator) (Figure 4B). To record from two locations within the chamber simultaneously, two Neuropixels probes are adhered to each other. Double-sided tape (Scotch, Double-Coated Tape) is placed on either side of a thin piece of plastic which serves as a spacer. To achieve the desired distance between the probes, spacers are made with different widths (used 1 mm or 1.9 mm). The spacer is sandwiched between the tail ends of the two probes, one of which has a holder attached (Figure 4C-E). Polyimide tape is then used to ensure the probes stay together and to keep the connector wires together.

**Figure 4.**
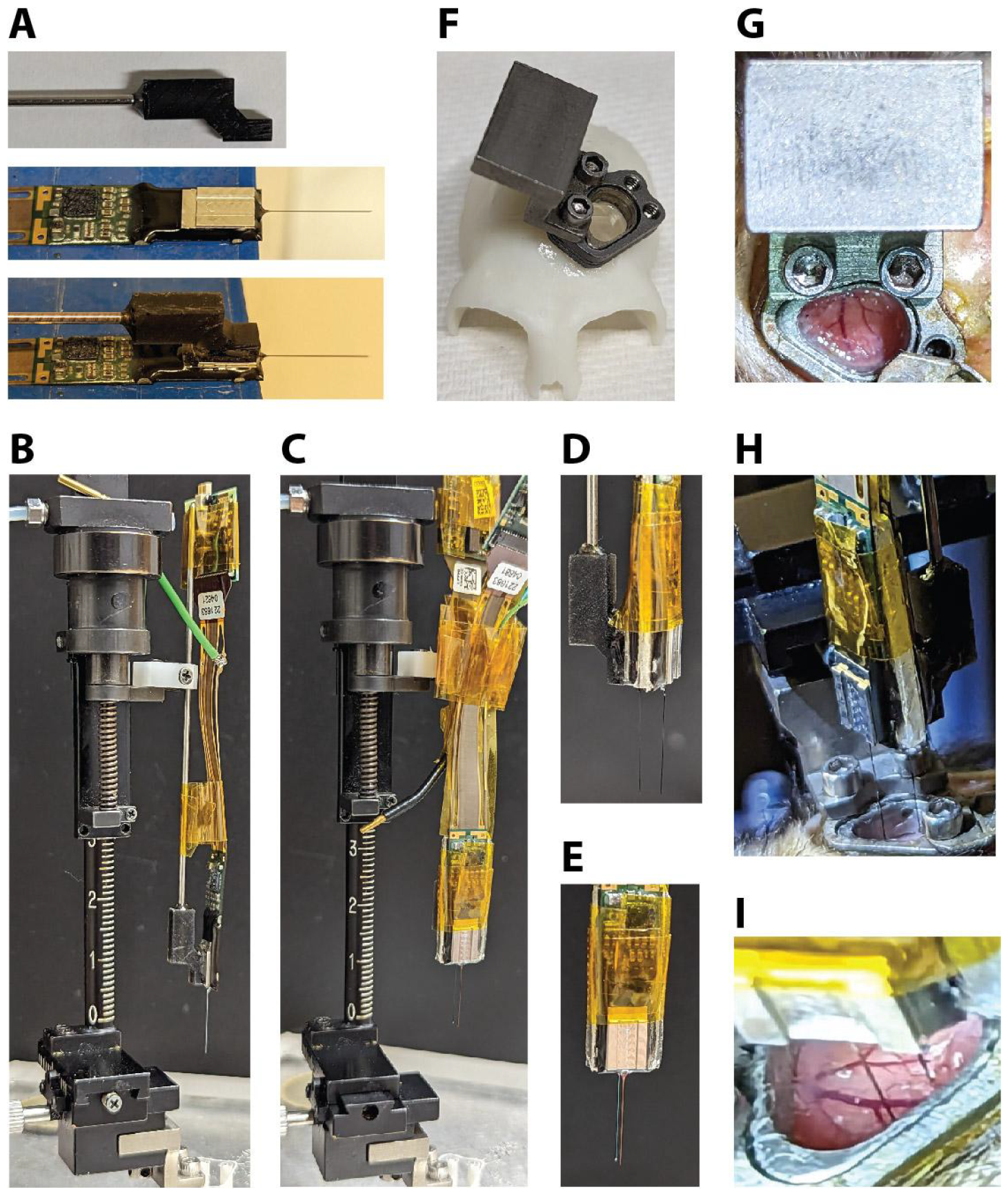
Neuropixels recording setup. A) A custom holder with a metal rod is glued to the base of the Neuropixels probe. Ground and reference wires are also attached. B) The rod is held by the clamp of the microdrive. Tape is used to control the cable and headstage. C) Example of double probe setup attached to microdrive (1.0 mm spacer). D) Side view and E) front view of double probe setup. F) 3D-printed skull model made from a CT scan with a chamber and insert (AD attached) and a microdrive mount attached. G) Photo of the microdrive mount when it is attached to the chamber. H) Photo of the double probe setup during the process of lowering the probes (1.9 mm spacer). I) Close-up view of the two probes while inserted in the cortex. Note the visible blood vessels that could be avoided during probe insertion.

Before attaching the microdrive, the outside of the chamber and the cap are thoroughly disinfected and the cap is then removed using sterile tools. A microdrive mount made from titanium is then attached to the side of the chamber using the same screw holes as the cap (Figure 4F,G). The microdrive holding the probe/s is then attached to the microdrive mount (Figure 4H, also see Figure 4B,C). Once the recording location has been determined using the x-y stage of the microdrive, the probes are then slowly lowered into the brain through the AD (Figure 4H,I).

After lowering the probe and allowing it to settle for approximately 30 minutes, recordings are then conducted for approximately 2 hours while the marmoset performs a variety of tasks, including viewing and saccading to a perceptual illusion (Dotson et.al. 2023), visual response field mapping, and full-screen flash sessions used to calculate the current source density (CSD). CSD plots are used to estimate laminar compartments based on the patterns of current sinks and sources and is calculated by taking the second spatial derivative of the average local field potential (LFP) response to a full-screen flash (Mitzdorf & Singer, 1978; Mitzdorf, 1985; Franken & Reynolds, 2021; Davis, et. al., 2023). We found that there was a common pattern of sources and sinks, with the earliest sink occurring near the middle of the cortex indicative of the “input” layer (layer 4) of the cortical column (Figure 5A). Unit activity was reliably found to occur within a millimeter above and below the estimated input layer (Figure 5B).

**Figure 5.**
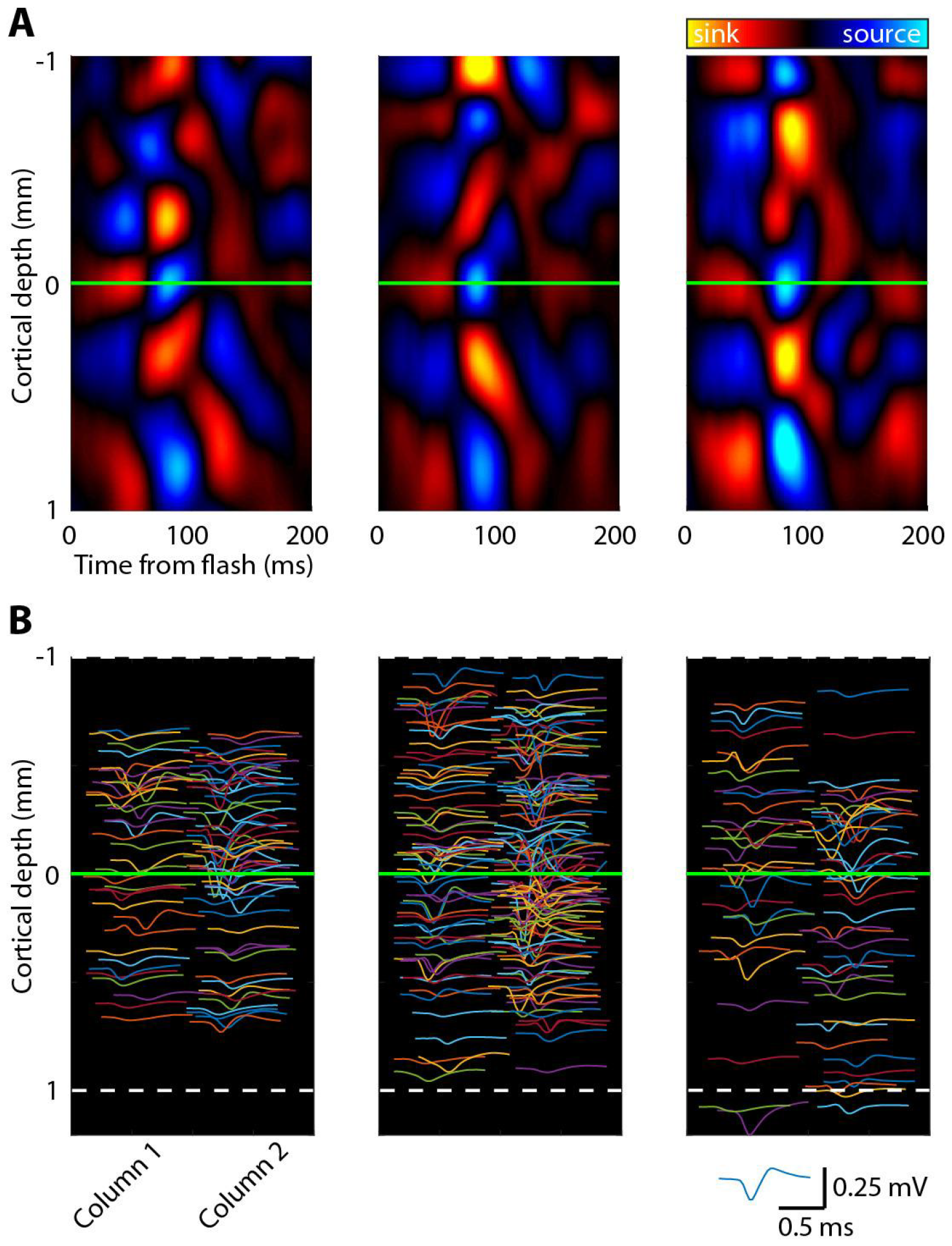
Current source density (CSD) can be used to identify the input layer of the cortex. Examples of CSD plots and spiking data from 3 recording sessions using Neuropixels probes. A) CSD plots (arb. units) with the input layer highlighted (green horizontal line). B) Average waveforms for each detected unit are shown at the relative depth and relative location (column 1 or column 2) on the probe (jittered for visibility). Number of isolated units: left (n = 215), middle (n = 255), and right (n = 144).

To reconstruct the location of the recording chamber with respect to the cortex, we first built a 3D model of the skull made from a CT scan taken after the headpost and chamber were implanted (Figure 6A,B), and then aligned a 3D model of the chamber (Figure 6C-D). We then registered a separate 3D model of the skull (made from a CT scan taken before implanting), with a 3D model of the marmoset brain inside (Liu et. al., 2021; marmosetbrainmapping.org), to the implanted skull model. After digitally removing the skull we could estimate the location of each cortical area with respect to the chamber (Figure 6E). To estimate the location of the probe after each recording session, the microdrive is mounted on a pedestal and chamber identical to the ones used for recordings, and a picture is taken from below (Figure 6F). This provides the location of the probe with respect to the chamber. This information can be used to identify the location and cortical area the probe likely penetrated (Figure 6G).

**Figure 6.**
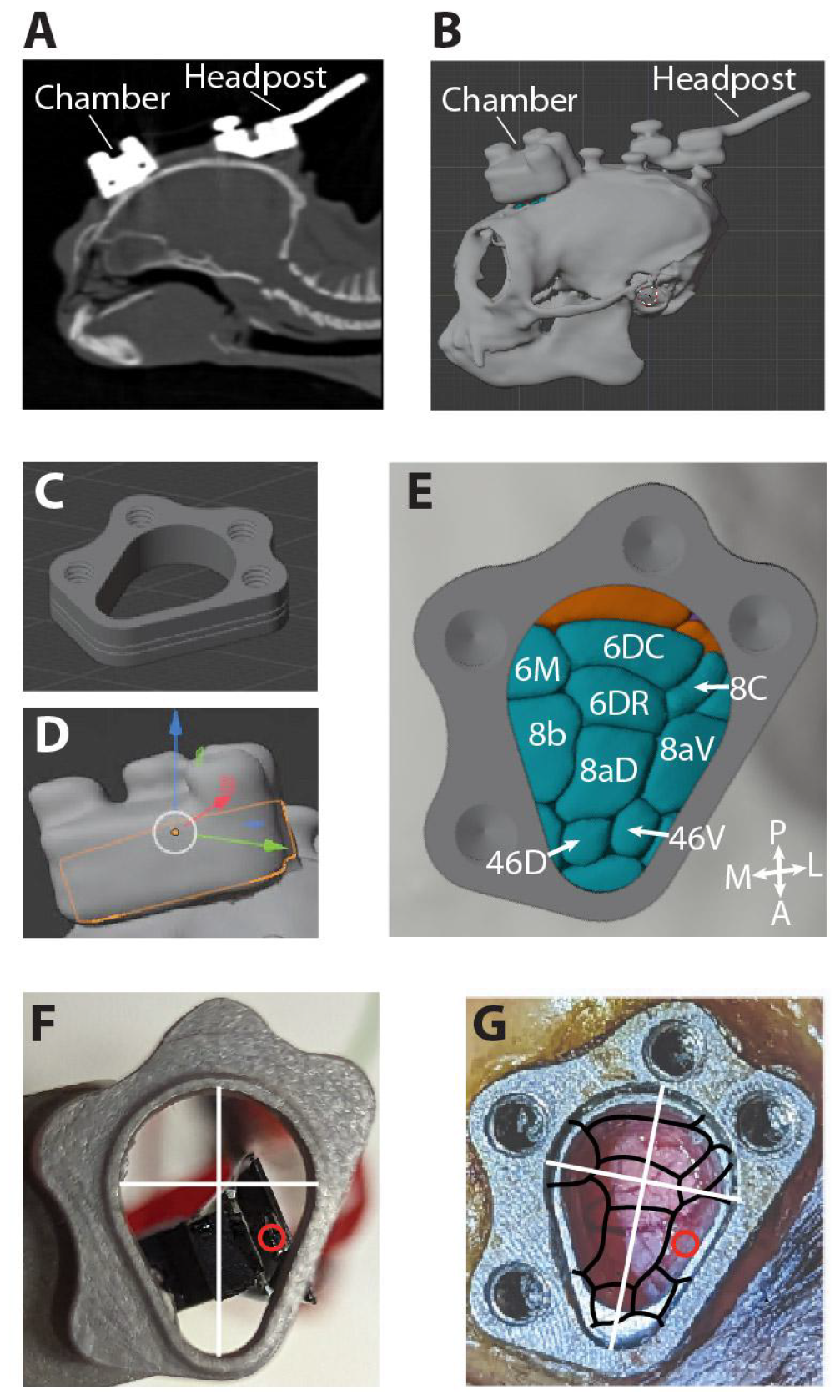
Reconstruction of recording locations. Recording locations were identified by combining 3D models and images taken of the probe and chamber. A) CT scan after head post and chamber implant. B) 3D-model of skull created from CT scan. C) 3D-model of chamber. D) Chamber model aligned to the chamber on the 3D-model of skull. The chamber model is transparent with an orange outline. E) Image through chamber of 3D-brain model (Liu et. al., 2021; marmosetbrainmapping.org). Anterior (A), posterior (P), medial (M), and lateral (L) directions are indicated in the bottom right corner. F-G) A separate chamber is used as a reference to determine the probe location relative to the chamber. After each recording, the microdrive is mounted on a pedestal and chamber identical to the ones used for recording (F). A picture is taken from below and then annotated. The chamber references (white lines) and probe shank location (red circle) are placed on top of the reconstructed map of areas to identify the recording location (G). This particular recording session is likely in area 8aV. Double probe recording sessions are reconstructed in the same manner.

## DISCUSSION

The shape of the marmoset cortex provides a unique opportunity to study laminar processing. Here we demonstrate an efficient method for acute Neuropixels recordings in the behaving marmoset. The AD recording system enables repeatable acute recordings with high optical visibility to avoid damaging blood vessels and a low risk of infection over an extended period of time. This technique has been successfully combined with marmosets performing a variety of complex behavioral tasks (e.g., Dotson et. al., 2023), to enable an exceptional ability to study laminar processing in a primate.

While the method presented here offers numerous benefits, there are also limitations including having to perform routine removal of granulation tissue. In some instances, it was not feasible to place the AD directly on the pia, leaving a small fluid-filled gap. This gap did not prove to be a problem for probe insertion but it may have shortened the window for recording by allowing tissue growth to more easily occur on the pial surface. Although the chambers used here were flat on the bottom, using form-fitting chambers (Dotson et. al., 2015. Dotson et. al., 2017; Jendritza et. al., 2023) and inserts may help to achieve a smaller gap and thus extend recording periods. The technique for holding probes and recording from multiple probes could also be improved. Future work could optimize the current method, which relies on epoxy and adhesive tape, by designing custom 3D-printed or machined parts that precisely align and secure the probes in position.

With some modifications, this method could be integrated with optical imaging (Pattadkal et. al., 2022; Song et. al., 2022) and optogenetics (Macdougall et al., 2016; Komatsu et al., 2017; Ebina et al., 2019; Jendritza et al. 2023; Obara et. al., 2023) in the head-fixed marmoset. Another avenue for future work will be to combine this technique for acute Neuropixels recordings with semi-chronic (Dotson et. al., 2015. Dotson et. al., 2017; Dotson et. al., 2018; Pomberger & Hage, 2019; Jendritza et al. 2023), and wireless recording methods (Courellis et. al., 2019; Dotson et. al., 2021; Mao et. al., 2021; Walker et. al., 2021; Wong et. al., 2023; Piza et. al., 2023) to understand the laminar mechanisms of visual motion and self-motion perception during natural movements in three dimensions. Chambers and semi-chronic microdrives capable of housing Neuropixels probes may be designed using CT scans to fit specific monkeys and target specific cortical regions (Dotson et. al., 2015. Dotson et. al., 2017; Jendritza et. al., 2023).

## MATERIALS AND METHODS

### Chamber implant procedure

For the PFC implant, the chamber is placed over the prefrontal cortex using the midline and the orbital arch as landmarks. First, an incision is made down the midline of the skull and the underlying tissue is removed from the bone. A thin layer of dental bonding agent (OptiBond Universal, Kerr) is then applied to the skull. Next, a craniotomy is made and several screws are placed around the perimeter. The chamber is then placed over the craniotomy and fixed in place using a fast-curing acrylic (Jet™ Acrylic, Lang Dental). Following the durotomy, the insert with the attached AD is lowered into the chamber. The durotomy does not necessarily have to be performed during the same surgery. For one of the PFC implants, the durotomy was performed several months after the chamber implant. With the occasional surgical removal of granulation tissue growth, acute Neuropixels recordings can be made for over a year. The chamber is rinsed with sterile saline every 5 days or less. Each time the chamber is opened the gasket and cap are replaced with sterile ones.

### Neural recording setup

Prior to neural recordings, marmosets are trained to freely enter custom-made marmoset chairs, which are then positioned in front of a calibrated and gamma corrected LCD monitor (ASUS VG248QE; 100 Hz refresh rate; 75 cd/m^2^ background luminance). Eye position is measured with a video-based eye tracker (ISCAN ELT-200, 500Hz sampling rate). Stimulus presentation and behavioral control are performed using MonkeyLogic (Asaad & Eskandar, 2008; Hwang et. al., 2019). Digital and analog signals are coordinated through National Instrument DAQ cards (NI PCI6621) and BNC breakout boxes (NI BNC2090A). Neuropixel recordings are acquired using the PXIe Acquisition Module (Neuropixels), and SpikeGLX acquisition software. A synchronization pulse is sent to both the PXIe Acquisition Module and the National Instrument DAQ card. After lowering the probe and allowing it to settle for 30 minutes, recordings are then conducted for ∼2 hours. Spike sorting is performed using Kilosort 2.0 and Phy (Pachitariu et. al., 2016; https://github.com/cortex-lab/phy). The chamber is rinsed with sterile saline before and after each recording.

## ACKNOWLEDGEMENTS

We thank C. Williams for help with the animals, S. Barry for machine shop work and helpful discussions, and T. Franken for helpful discussions. Funding was provided by the Fiona and Sanjay Jha Chair in Neuroscience (J.H.R), George E. Hewitt Foundation for Medical Research (P.J), R01-EY028723 (J.H.R., Z.W.D., N.M.D.) with early foundational support provided through grants T32 EY020503-06, P30 EY019005.

## Notes

### Competing Interest Statement

The authors have declared no competing interest.

